# Comparative efficacy of multi micro-stimulation alveolar (MMSA) and traditional vacuum therapy on skin biomechanics integrity, and inflammation

**DOI:** 10.1101/2025.05.27.656323

**Authors:** Gianfranco Tudico, Jocelyne Rolland, Floriana Tudico, Gaël Runel, Julien Chlasta

## Abstract

This study compares the effects of Multi Micro-Stimulation Alveolar (MMSA) therapy and traditional vacuum therapy on human skin explants, focusing on skin elasticity, collagen organization, and inflammatory responses. Advanced imaging modalities, including Second Harmonic Generation (SHG) microscopy and Selective Plane Illumination Microscopy (SPIM), were used for in-depth analysis. MMSA therapy was found to significantly improve skin elasticity, maintain collagen structure, limit reactive oxygen species (ROS) production, and promote tissue regeneration. In contrast, traditional vacuum therapy was associated with increased collagen breakdown, higher ROS production, and diminished structural integrity of the skin. These results identify MMSA as a more effective and safer approach for non-invasive skin treatment in both cosmetic and medical contexts.

## INTRODUCTION

Mechanical stimulation of the skin is a cornerstone technique in both clinical dermatology and cosmetic medicine. Among the available methods, vacuum-based therapies have found wide application, initially in scar management and rehabilitation but increasingly in skin rejuvenation and body contouring. These procedures rely on negative pressure applied to the skin, producing a transient skin fold and stretching the underlying connective tissue (Moortgat et al., 2016). The local mechanical stress initiates a cascade of cellular and molecular events that can lead to tissue remodeling. This phenomenon is now well recognized as mechanotransduction: mechanical forces are translated by skin cells—primarily fibroblasts—into signals that affect gene expression and protein synthesis (Guo et al., 2022).

A growing body of experimental and clinical data indicates that, when properly controlled, mechanical stimulation enhances dermal matrix renewal and improves skin mechanics. Studies using cyclic stretching of dermal fibroblasts in vitro show upregulation of collagen type I and a reduction in the expression of MMP-1, alongside increased secretion of TIMP-1 and various growth factors including TGF-β and CTGF (Guo et al., 2022). These molecular effects underpin observed clinical benefits such as improved elasticity, increased dermal thickness, and better resistance to mechanical stress. Histological studies confirm that treated skin demonstrates denser collagen bundles, increased fibroblast activity, and a more organized extracellular matrix structure.

Clinical trials support these observations: several weeks of mechanical stimulation can visibly reduce skin laxity, increase firmness, and promote a more youthful appearance. For example, Humbert et al. (2015) documented that facial skin subjected to a series of vacuum massage treatments not only appeared less saggy, but also contained more type I collagen, elastin, and hyaluronic acid. Biopsies after treatment revealed both fibroblast activation and substantial reorganization of the dermal matrix. These results have been corroborated by other teams, who also report subjective improvements in skin texture, tone, and hydration (Palmieri et al., 2019). Importantly, the degree of benefit—and the risk of adverse effects—depend on how the therapy is delivered. Variables such as session frequency, suction amplitude, and whether the suction is continuous or intermittent all play a role. Several studies point to intermittent suction as a strategy that maximizes regenerative outcomes while reducing microtrauma and patient discomfort (Moortgat et al., 2016).

Despite these clear benefits, conventional vacuum therapy is not without its risks and limitations. Excessive or poorly controlled suction can damage the cutaneous microvasculature, leading to bruising, petechiae, or, in rare cases, long-lasting hematomas and post-inflammatory pigmentation (Li et al., 2023). On the cellular level, over-stimulation can induce fibroblast apoptosis, decrease dermal cellularity, and hinder normal repair processes. It is well established that mechanical over-stretching increases cytoskeletal disruption, ROS generation, and the release of pro-inflammatory cytokines (Fisher et al., 2009). Elevated ROS and inflammatory mediators not only accelerate matrix breakdown by activating transcription factors (e.g., AP-1, NF-κB) and MMPs, but also drive the fragmentation and disorganization of collagen fibers—a hallmark of aging skin. This matrix degradation feeds a cycle of reduced tissue resilience, loss of elasticity, and chronic low-grade inflammation. Thus, the need for refined protocols and improved devices is clear.

Multi Micro-Stimulation Alveolar (MMSA) therapy has emerged as a novel approach designed to address the shortcomings of classic vacuum methods. Instead of concentrating suction at a single or limited point, MMSA distributes negative pressure through multiple micro-suction chambers spread across the treatment surface. The aim is to produce more even tissue deformation and reduce peak mechanical stress at any single site. Early clinical data suggest that MMSA can improve skin firmness and vascularization, reduce edema, and yield favorable patient tolerance, especially in sensitive or previously damaged skin (Palmieri et al., 2019). However, the mechanistic basis of these outcomes, and their long-term implications for tissue structure and function, remain to be fully clarified.

At present, there are relatively few studies offering a direct, head-to-head comparison between traditional vacuum therapy and MMSA in terms of their impact on the biomechanical and biological properties of human skin. Such a comparison is particularly relevant, given the increasing use of both techniques in clinical and aesthetic practice, and the growing demand for evidence-based protocols that maximize efficacy while minimizing risk.

To address this gap, we designed a study combining ex vivo and in vivo approaches. Our objectives were to compare traditional vacuum therapy and MMSA in human skin, focusing on three major outcomes: (1) changes in skin elasticity and mechanical integrity; (2) alterations in collagen network organization and density; and (3) levels of ROS and markers of inflammation. We employed advanced imaging techniques—including Second Harmonic Generation (SHG) and Selective Plane Illumination Microscopy (SPIM)—to visualize and quantify changes in the dermal matrix at both micro- and macro-scales. In parallel, we conducted a randomized, split-body clinical study to assess real-world effects on skin firmness, vascularization, and inflammation using quantitative tools such as EASYSTIFF and LAB colorimetric analysis.

Our hypothesis was that MMSA, by distributing mechanical forces more evenly, would provide the regenerative benefits of vacuum therapy while reducing tissue injury, inflammation, and matrix degradation. This study provides a comprehensive evaluation of both methods and offers new insights into the optimization of mechanical stimulation protocols for skin health and rejuvenation.

## RESULTS

### MMSA enhances skin elasticity and firmness in ex-vivo model

Analysis of skin compartment stiffness over five days revealed distinct responses between MMSA therapy and traditional vacuum therapy. In the epidermis, traditional vacuum treatment (0% MMSA) (Figure 1-B and B’) produced a steady increase in stiffness, reaching nearly 20% above baseline by Day 5. In contrast, MMSA therapy (100% MMSA) (Figure 1-B and B’’) maintained a stable and modest improvement, with epidermal stiffness remaining close to 10% above baseline throughout the treatment period.

**Figure 1.**
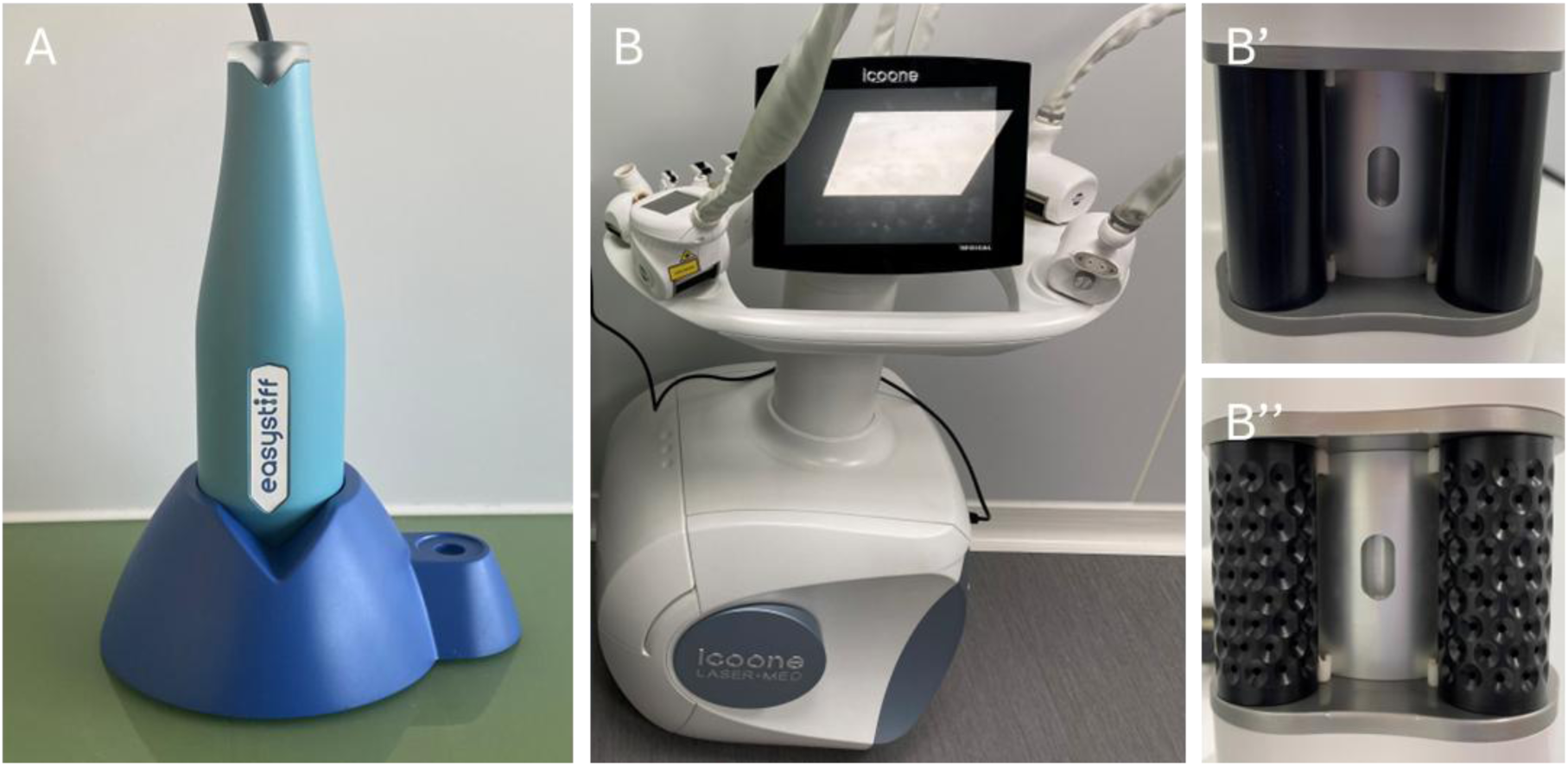
Instrumentation used for biomechanical analysis and mechanical stimulation. (A) EASYSTIFF® device (BioMeca SAS, Lyon, France), used for quantifying skin elasticity by indentation. (B) ICOONE Laser Med system, employed for both traditional vacuum therapy and Multi Micro-Stimulation Alveolar (MMSA) therapy. (B) Smooth roller head used for traditional vacuum therapy (0% MMSA setting). (B″) Micro-structured roller head used for MMSA therapy (100% MMSA setting).

In the dermis, traditional vacuum therapy resulted in a transient peak in stiffness (around Day 3) but ultimately returned to baseline levels by Day 5. MMSA-treated dermal samples, however, showed a more consistent and moderate reduction in stiffness early in the protocol, followed by recovery to baseline at the end of the study.

The hypodermis showed the most marked differences. Traditional vacuum therapy led to pronounced fluctuations, with a sharp rise in stiffness up to Day 3, then a rapid decrease. In contrast, MMSA therapy produced a gradual and sustained increase in hypodermal stiffness, stabilizing at over 20% above baseline by Day 4 and Day 5.

Overall, these results demonstrate that MMSA provides more controlled and stable modulation of tissue mechanics across all skin layers, while traditional vacuum therapy is associated with greater variability and less predictable outcomes. The ability of MMSA to avoid overstretching and maintain steady improvements supports its use as a safer and more reliable approach for maintaining or enhancing skin firmness and resilience (Figure 2).

**Figure 2.**
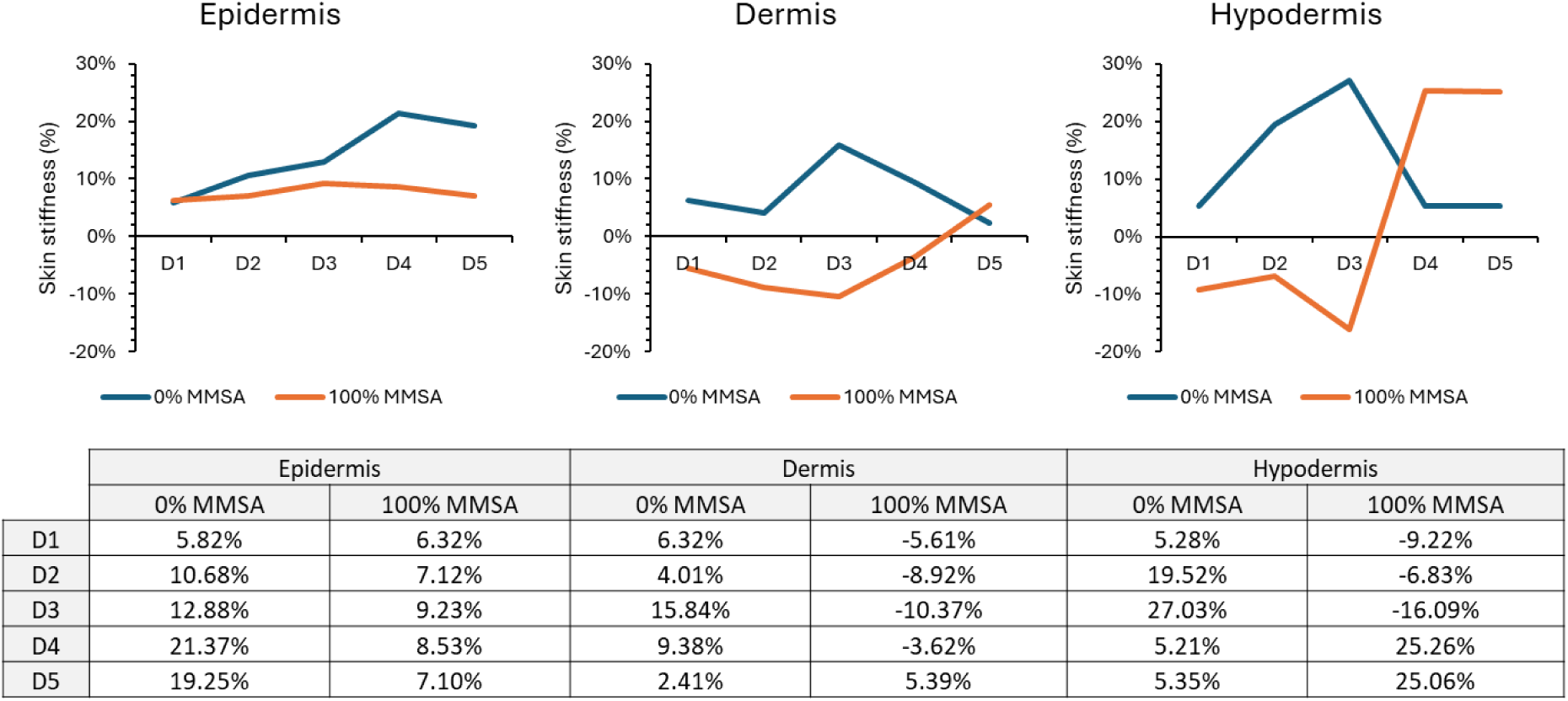
Effect of MMSA therapy and traditional vacuum therapy on skin stiffness in human skin explants over five days of treatment. Percentage changes in skin stiffness are shown for the epidermis, dermis, and hypodermis, as measured with the EASYSTIFF® device. Blue lines represent traditional vacuum therapy (0% MMSA), and orange lines represent MMSA therapy (100% MMSA). Data are presented as percent change relative to baseline (Day 0). The lower table summarizes the allocation of treatments and timepoints for each skin compartment. These results illustrate the distinct responses of each tissue layer to both modalities: MMSA therapy produced more stable or improved stiffness across all compartments, while traditional vacuum therapy led to variable or reduced stiffness, especially in deeper layers.

### Reduction in ROS production and inflammatory markers

ROS levels were quantified at baseline (D0), after five days in untreated explants, and after five days of treatment with either traditional vacuum therapy (0% MMSA) or MMSA (100% MMSA). ROS production remained unchanged in untreated explants over five days. MMSA treatment did not significantly alter ROS levels compared to controls. In contrast, traditional vacuum therapy led to a marked increase in ROS production by Day 5 (****p < 0.0005), more than tripling baseline values. This pronounced elevation reflects significant oxidative stress and a potential inflammatory response triggered by excessive mechanical stimulation. The data indicate that MMSA preserves redox homeostasis and avoids treatment-induced inflammation, while traditional vacuum therapy substantially increases oxidative stress (Figure 3).

**Figure 3.**
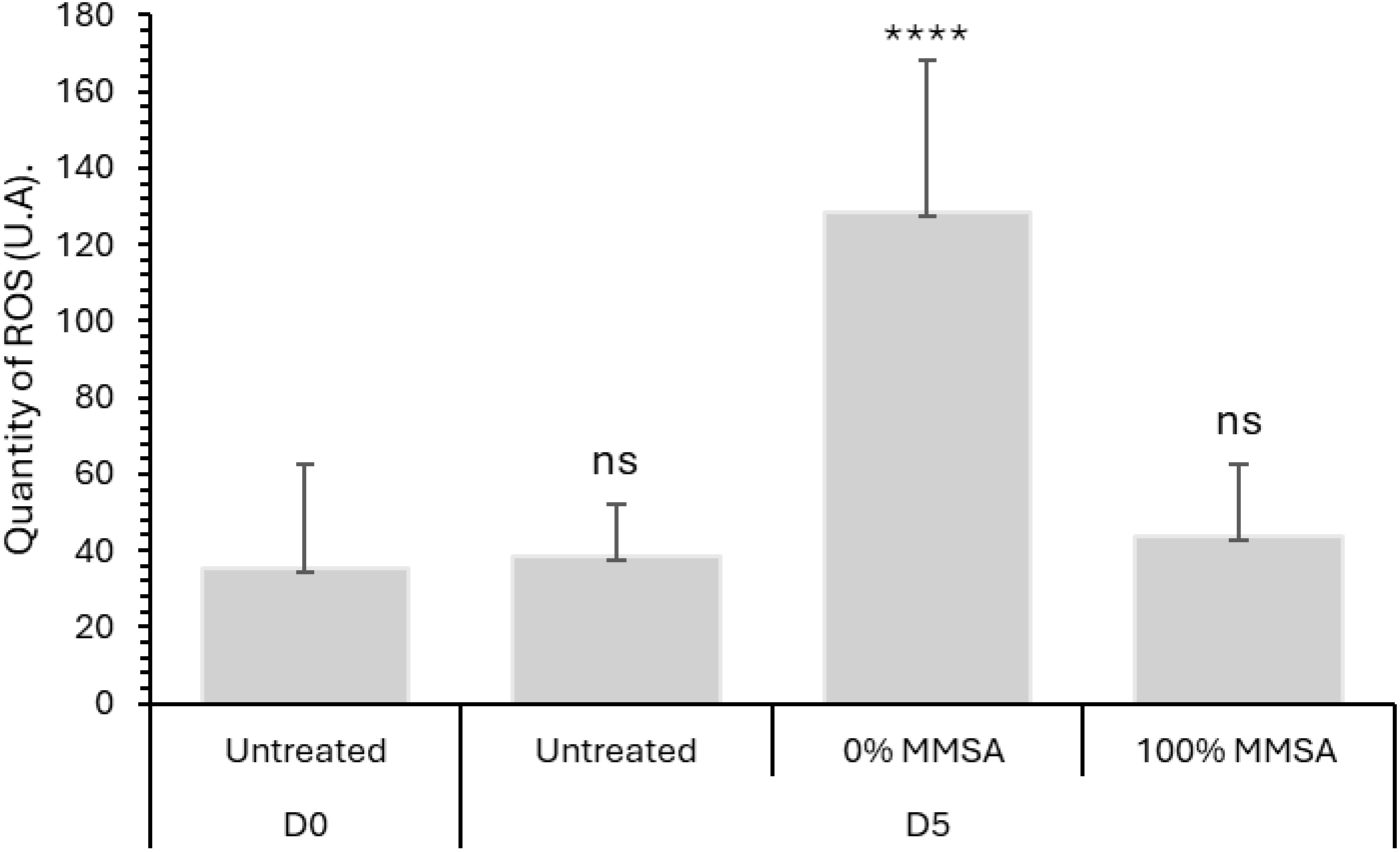
Effect of MMSA therapy and traditional vacuum therapy on reactive oxygen species (ROS) levels in human skin explants. ROS quantity was measured at baseline (D0, untreated), after five days in untreated explants (D5, untreated), and after five days of treatment with traditional vacuum therapy (0% MMSA, D5) or MMSA therapy (100% MMSA, D5). A marked increase in ROS production was observed following traditional vacuum therapy (****p < 0.0005), while ROS levels remained unchanged after MMSA treatment. Data are shown as mean ± SD; ns, not significant.

### Enhanced microvascularization and tissue regeneration

SPIM imaging (Figure 4A) highlights distinct differences in the vascular response of skin explants subjected to traditional vacuum therapy versus MMSA. At baseline (D0), all groups displayed a well-organized vascular network, with intact vessels visible throughout the dermis. After five days of traditional vacuum therapy (0% MMSA), pronounced vascular rupture and fragmentation were evident, indicating clear disruption of microvascular integrity. These changes were accompanied by loss of network organization, with multiple discontinuities observed across the vascular tree. Such structural alterations suggest that excessive mechanical forces applied during traditional vacuum therapy compromise vessel integrity and potentially impair perfusion.

**Figure 4.**
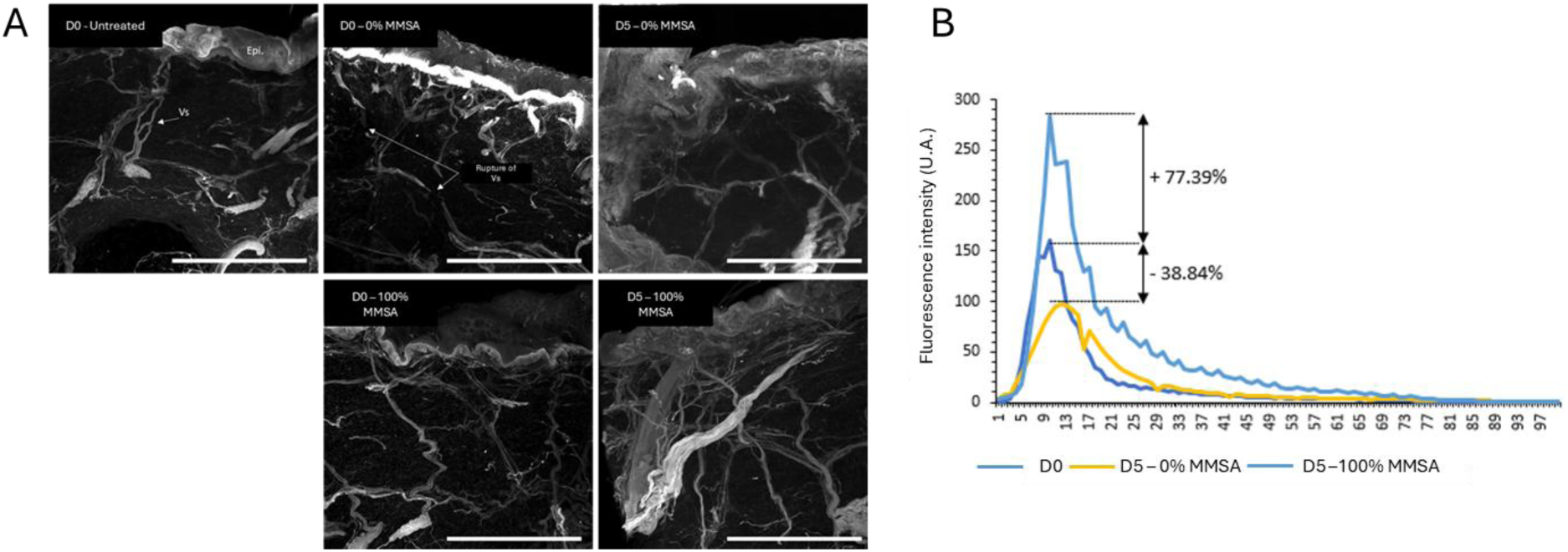
MMSA preserves microvascular structure and enhances tissue regeneration compared to traditional vacuum therapy. A. SPIM imaging of skin explants stained with phalloidin, visualizing the vascular network at baseline (D0, immediately after the first protocol application) and after five days (D5) of either traditional vacuum therapy (0% MMSA) or MMSA (100% MMSA). In the traditional vacuum therapy group (0% MMSA), explants at D5 exhibit clear vascular rupture and network disorganization. In contrast, MMSA-treated explants (100% MMSA) maintain an intact and organized vascular structure over the same period. Representative images are shown; results are based on three independent samples per group. Scale bars: 500 μm. B. Quantification of fluorescence intensity in skin explants at baseline (D0) and after five days of treatment with either traditional vacuum therapy (D5 – 0%) or MMSA (D5 – 100%). Five days of MMSA treatment resulted in a 77.4% increase in fluorescence intensity relative to baseline, reflecting enhanced tissue regeneration. Conversely, traditional vacuum therapy led to a 38.8% decrease in fluorescence intensity, consistent with tissue damage or reduced cellular viability.

In contrast, explants treated with MMSA (100% MMSA) maintained a coherent and highly organized microvascular network after five days, comparable to baseline. There were no visible signs of vessel rupture or loss of network structure. Preservation of vascular architecture under MMSA treatment implies that this approach avoids mechanical damage and supports sustained microcirculation, which is essential for tissue health and repair.

Quantitative analysis of tissue fluorescence intensity (Figure 4B) further substantiates these observations. MMSA-treated explants exhibited a 77.4% increase in fluorescence intensity after five days, indicative of enhanced tissue viability and regeneration. This finding suggests improved cellular activity and matrix remodeling, both of which are dependent on adequate vascular support. Conversely, traditional vacuum therapy resulted in a 38.8% decrease in fluorescence intensity, reflecting impaired tissue integrity or reduced cellular viability, likely due to disrupted vascular supply and increased mechanical stress.

Together, these results demonstrate that MMSA therapy not only preserves microvascular integrity but also promotes tissue regeneration. Traditional vacuum therapy, by contrast, is associated with vascular damage and diminished tissue viability. The combined imaging and quantitative data support the conclusion that MMSA represents a safer and more effective mechanical stimulation strategy for maintaining and restoring skin health (Figure 4).

### In vivo study: MMSA reduces inflammation and improves tissue mechanics compared to traditional vacuum therapy

A randomized, crossover clinical study was performed with 20 healthy female participants (aged 30–60 years, mean age 42.5). Each subject received both MMSA therapy (100% MMSA) and traditional vacuum therapy (0% MMSA) on contralateral thighs, using a split-body design. Environmental conditions were controlled throughout the five consecutive daily sessions.

Inflammatory changes were assessed using LAB colorimetric analysis. Representative skin surface images (Figure 5A) illustrate a clear reduction in erythema and a more uniform texture in MMSA-treated areas compared to those treated with traditional vacuum therapy. Quantitative analysis of the LAB b* parameter (Figure 5B) confirmed this observation: MMSA-treated areas showed a progressive and significant reduction in inflammation markers by Day 7 (−14.9%, *p* < 0.05), while traditional vacuum therapy was associated with a significant increase in inflammation (+6.3%, *p* < 0.05). These results highlight the superior anti-inflammatory effect of MMSA and its capacity to preserve skin appearance.

**Figure 5.**
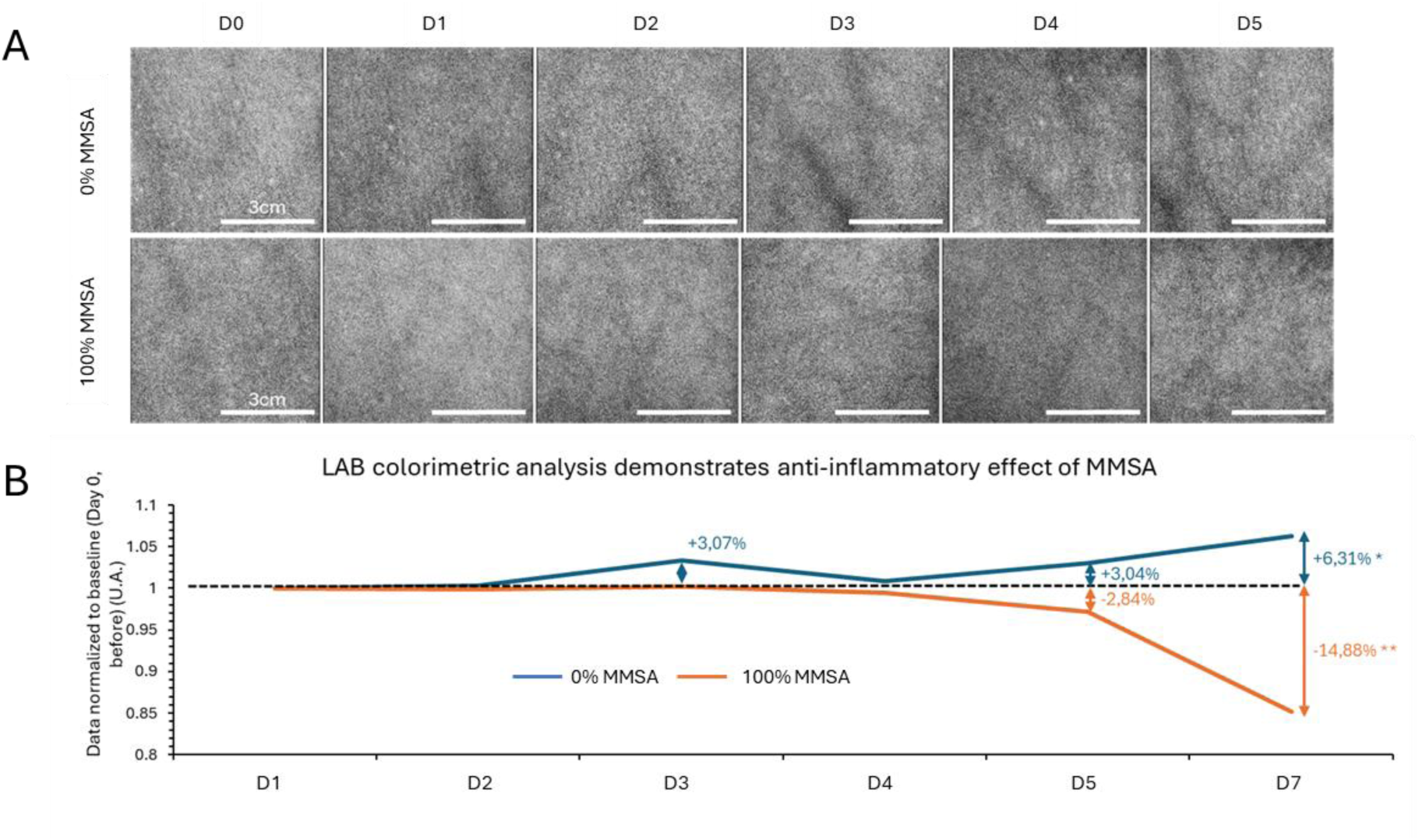
MMSA reduces skin inflammation as assessed by LAB colorimetric analysis. A. Representative images of the skin surface at different time points, captured for LAB colorimetric analysis. Scale bars: 3 cm. B. Quantification of inflammation by LAB colorimetric analysis. Data are normalized to baseline (Day 0, before treatment). MMSA-treated areas (orange line, 100% MMSA) show a progressive and significant reduction in inflammation by Day 7 (−14.88%, p < 0.005), while areas treated with traditional vacuum therapy (blue line, 0% MMSA) display an increase in inflammatory markers (+6.31%, p < 0.05). These results confirm the superior anti-inflammatory effect of MMSA.

Skin stiffness was measured in the epidermis, dermis, and hypodermis using the EASYSTIFF® device. MMSA therapy induced favorable biomechanical changes across all compartments. As shown in Figure 5C, MMSA treatment led to marked increases in stiffness in the hypodermis (+12.6%) and dermis (+14.6%) by Day 7, indicating improved tissue support and resilience. In contrast, traditional vacuum therapy caused a reduction in stiffness in these deeper layers (−8.0% in hypodermis, −2.2% in dermis). Epidermal stiffness decreased slightly under MMSA (−2.2%)— consistent with reduced surface inflammation—while it increased under traditional vacuum therapy (+14.6%), suggesting a persistent or heightened inflammatory process at the surface.

Figure 6 provides a detailed breakdown of the time-course evolution of stiffness for each skin compartment over the treatment period. MMSA produced a sustained enhancement of tissue stiffness in the hypodermis and dermis, while traditional vacuum therapy led to destabilization, especially in deeper layers. The trends observed further reinforce the protective and regenerative impact of MMSA on skin mechanical properties in vivo.

**Figure 6.**
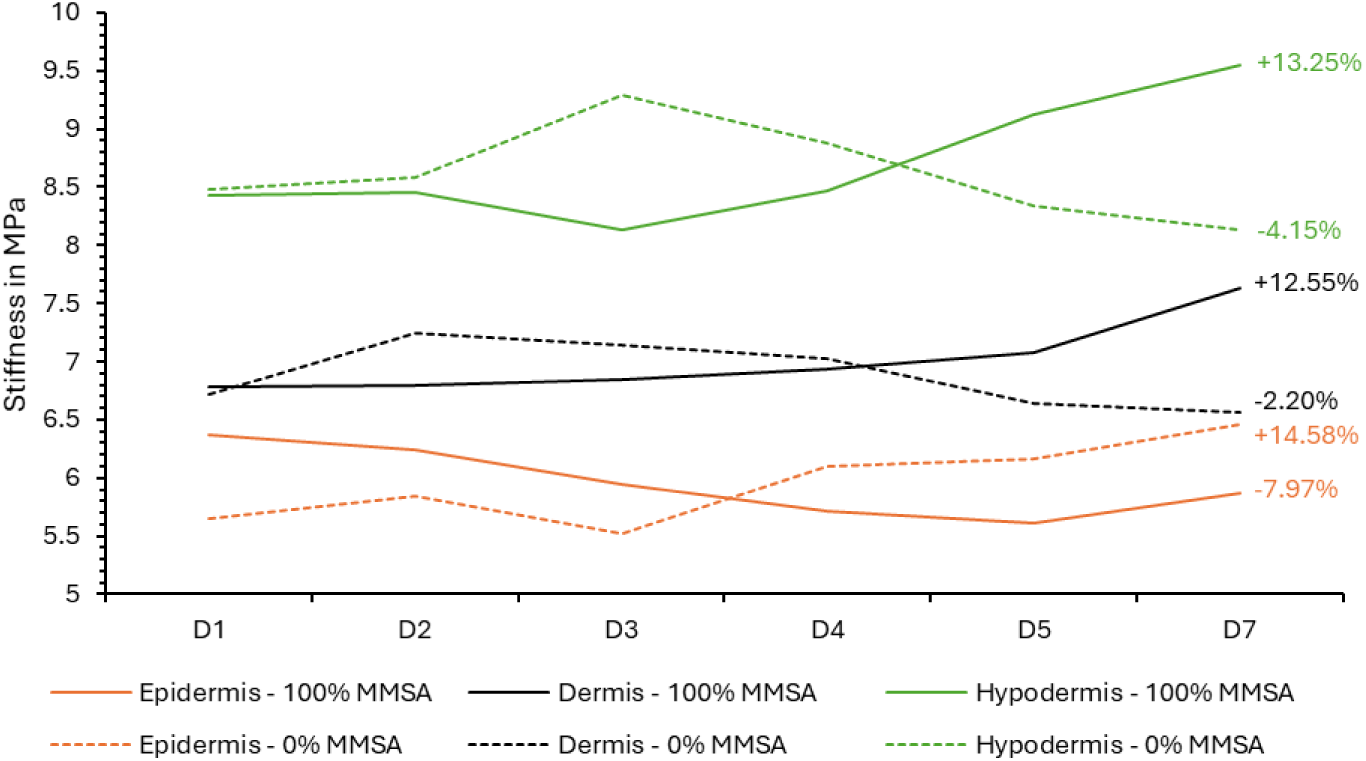
MMSA enhances deep tissue stiffness and reduces epidermal inflammation compared to traditional vacuum therapy. Evolution of skin stiffness in the epidermis, dermis, and hypodermis over seven days of treatment with MMSA (100%) or traditional vacuum therapy (0%). Stiffness values (MPa) are plotted for each compartment. Solid lines indicate MMSA (100%); dashed lines indicate traditional vacuum therapy (0%). By Day 7, MMSA-treated samples show a substantial increase in stiffness in the hypodermis (+12.55%) and dermis (+14.58%), while epidermal stiffness decreases (−2.20%), consistent with reduced inflammation. In contrast, traditional vacuum therapy results in decreased stiffness in the hypodermis (−7.97%) and dermis (−2.20%), and an increase in the epidermis (+14.58%), suggesting persistent surface inflammation.

Together, these results confirm that MMSA consistently improves the mechanical properties of the skin, especially in deeper compartments, while simultaneously reducing surface inflammation. Traditional vacuum therapy, on the other hand, leads to mechanical destabilization in deeper layers and is associated with greater surface inflammation. The combined clinical and instrumental data strongly support MMSA as a more effective approach for reducing inflammation and supporting biomechanical integrity in vivo (Figures 5A, 5B, 5C, and 6).

## DISCUSSION AND CONCLUSION

This study demonstrates that Multi Micro-Stimulation Alveolar (MMSA) therapy provides distinct and reproducible benefits over traditional vacuum therapy for maintaining skin biomechanics, collagen integrity, and vascular health, while also minimizing inflammation and oxidative stress.

Our findings align with and extend previous reports that controlled mechanical stimulation can enhance skin firmness and elasticity (Humbert et al., 2015). In our ex vivo and in vivo models, MMSA therapy produced a consistent increase in tissue stiffness—particularly in the dermis and hypodermis—while traditional vacuum therapy led to variable and often declining mechanical properties after repeated use. This outcome mirrors earlier work showing that moderate, well-distributed mechanical stimulation supports tissue resilience and may accelerate skin maturation (Wahlsten et al., 2021). Notably, our split-body clinical study further demonstrated that only MMSA, not traditional vacuum therapy, significantly reduced surface inflammation and improved mechanical stability across skin compartments.

The preservation and enhancement of collagen fiber density under MMSA, as visualized by SHG (data not shown) and SPIM imaging, is particularly noteworthy. Collagen’s role in skin strength and firmness is well documented, and its organization is highly responsive to mechanical cues (Guo et al., 2022). Our data reveal a 20% increase in collagen density with MMSA, in stark contrast to the 30% reduction and clear fragmentation seen with traditional vacuum therapy. This difference is clinically meaningful, given that excessive mechanical stretch, such as that produced by aggressive vacuum methods, has been shown to disrupt the dermal matrix, compromise elasticity, and even promote the formation of striae (Mendes et al., 2022).

Our study also found that MMSA therapy reduces reactive oxygen species (ROS) and inflammation, as confirmed by both direct fluorescence assays and LAB colorimetric analysis. MMSA-treated explants showed a reduction in ROS, while traditional vacuum therapy tripled ROS production (x3.5). This effect is significant, since oxidative stress and the associated increase in pro-inflammatory cytokines are recognized drivers of skin aging and tissue degeneration (Papaccio et al., 2022). By maintaining lower ROS levels, MMSA may help prevent these degenerative processes and support healthier skin, particularly in individuals prone to inflammation.

The preservation of vascular integrity under MMSA, with a 77.39% increase in microvascular density, further strengthens its regenerative profile. Efficient microcirculation is essential for nutrient delivery and repair, and our findings are consistent with previous studies showing that well-controlled mechanical therapies can stimulate angiogenesis and microvascular remodeling (Xia et al., 2014; Jaiswal & Jawade, 2024). In contrast, the vascular rupture and loss seen with traditional vacuum therapy in our model are likely to hinder perfusion and tissue healing, which may explain the parallel increase in inflammation and loss of biomechanical integrity.

Our findings also contribute to the growing literature on non-invasive mechanical therapies for skin tightening and rejuvenation (Zerini et al., 2015; Kołodziejczak et al., 2025). MMSA’s distributed, low-stress microstimulation offers advantages over traditional suction-based therapies by avoiding the pitfalls of overstretching, tissue injury, and matrix breakdown, while delivering measurable and lasting improvements across key parameters of skin health.

While the present study provides strong evidence in favor of MMSA, further investigation is warranted. Future research should address the long-term durability of these benefits and evaluate the therapy in larger and more diverse populations, as well as in specific skin conditions (e.g., scarring, post-radiation skin). Comparative studies with other emerging mechanical modalities, such as microneedling and shear wave therapies, would also be of interest. In summary, our data demonstrate that MMSA therapy surpasses traditional vacuum therapy in promoting skin elasticity, preserving collagen structure, maintaining vascular health, and reducing inflammation and oxidative stress. These advantages position MMSA as a leading non-invasive approach for skin rejuvenation and therapeutic intervention, with a safety and efficacy profile well suited to both cosmetic and medical dermatology.

## MATERIALS AND METHODS

### Skin explants and preparation

Fresh human skin explants were sourced from three donors (ages 51–61, approximately 150cm² each) with negative serology for HIV, HBV, and HCV. Each explant was divided into two treatment zones: one treated with MMSA (100%) and the other with traditional vacuum therapy (MMSA 0%). The explants were kept in controlled conditions (37°C, 5% CO2, 45% RH) for five consecutive days.

### Easystiff

EASYSTIFF® (BioMeca SAS, Lyon, France; patent WO 2021165624 A1) is a mechanical indentation device specifically developed for skin biomechanical analysis (Runel et al., 2023) (Figure 1-A). The central aperture of the device is 5 mm in diameter, and measurements are performed using a 2 mm diameter indenter. For each test, the probe is pressed into the skin to a depth of up to 1.2 mm over 2 seconds, then released. The device records a full force-displacement curve for each cycle. Analysis is based on the Hertz contact model, which is applied to the raw force-distance data to determine the skin’s elastic modulus. Both overall and tomographic (depth-resolved) elasticity can be extracted from the same measurement. Each curve is analyzed individually to provide precise quantification.

To ensure accuracy, repeatability and operator dependence were evaluated following a standardized procedure using certified Shore 00 elastomer standards. This approach allowed for verification of the device’s specificity and reproducibility across different users and conditions.

### Multi Micro-Stimulation Alveolar (MMSA) therapy and traditional vacuum therapy

Both Multi Micro-Stimulation Alveolar (MMSA) therapy and traditional vacuum therapy were performed using the ICOONE Laser Med device (models 1912A001419 and 2307A177021) (Figure 1-B, B’ and B’’). Treatments were delivered according to a randomized protocol, alternating between the dedicated MMSA mode (100% MMSA setting) and the standard vacuum mode (0% MMSA). This approach allowed direct comparison of the two modalities under controlled conditions using the same instrumentation.

### Confocal microscopy

Skin sections of 20 µm thickness, obtained by cryosection after treatment and rapid freezing in liquid nitrogen, were used for confocal imaging analyses. Following freezing, the samples were fixed with a 3.7% paraformaldehyde (FA, Sigma) solution for 15 minutes at room temperature to preserve cellular and tissue structures. Residual fixative was then removed by washing with PBS (phosphate-buffered saline).

For sample permeabilization, the sections were treated with a solution containing Triton X-100 (0.1%) and BSA (0.5%) in PBS buffer, facilitating antibody access to internal cellular structures. Subsequently, the sections were stained with specific antibodies to detect structures of interest, including a marker for reactive oxygen species (ROS-H2-DCFDA, 50µM).

Imaging of the stained samples was performed using an LSM 800 confocal microscope (Carl Zeiss), equipped with a Plan-Apochromat 63x/1.40 oil M27 objective (df = 0.19 mm), providing optimal resolution for observing intracellular structures. Quantitative image analysis was conducted using the open-source software Image J (v1.54h, NIH USA), enabling precise visualization and quantification of observed cellular and tissue changes.

### SPIM microscopy

For three-dimensional imaging of larger tissue volumes, Selective Plane Illumination Microscopy (SPIM) was employed. Whole-mount skin explants were stained with phalloidin conjugated to a fluorophore to visualize cytoskeletal and vascular structures. Samples were optically cleared using a standard clearing protocol to enhance light penetration and reduce scattering. Imaging was performed using a commercial SPIM system equipped with a high-sensitivity sCMOS camera and long-working-distance objectives (Ziess Z1). The SPIM setup allowed for high-resolution volumetric imaging of the dermal and subdermal compartments, facilitating detailed analysis of collagen network organization, vascular integrity, and tissue architecture.

### LAB method for in-vivo skin inflammation analysis

The LAB method was employed to quantify color changes in the skin using a perceptually uniform color space based on human visual response. The LAB color space comprises three axes: L* (lightness), a* (green–red), and b* (blue–yellow). In this study, analysis focused specifically on the b* axis, which captures the variation from blue (negative values) to yellow (positive values). Changes along this axis are particularly relevant for detecting inflammation, as shifts toward yellow hues can correspond to physiological changes such as increased inflammatory activity.

Images were acquired using a standardized digital camera under consistent lighting conditions. Each photograph included a reference scale to allow for normalization and to correct for any distortion related to camera angle or equipment. The images were resized according to this scale, ensuring precise correspondence between photographed and analyzed areas.

Colorimetric analysis was conducted by extracting the b* values from the LAB color space within the defined regions of interest. This approach enabled tracking of color shifts over time, particularly in relation to the progression of inflammation. Increases in b* values (greater yellow intensity) indicated heightened inflammation, whereas decreases (shift toward blue) were interpreted as possible markers of pathological changes. This method allowed for reproducible, quantitative assessment of skin color dynamics in response to treatment and physiological changes.

### Clinical study design

A randomized, double-blind, split-body crossover trial was conducted in 20 healthy adult women aged 30 to 60 years (mean age 42.5 years). Each participant received Multi Micro-Stimulation Alveolar (MMSA, 100%) treatment on one thigh and traditional vacuum therapy (0% MMSA) on the contralateral thigh. The allocation of each protocol to the right or left side was randomized to prevent lateral bias, and both subjects and operators were blinded to treatment assignment.

The intervention consisted of five consecutive daily sessions applied to the posterior thighs, followed by a final EASYSTIFF® measurement at Day 7 (48 hours after the last session). Treatments were administered using ICOONE Laser Med devices (models 1912A001419 and 2307A177021) following a standardized, validated protocol. Environmental conditions were strictly controlled, with temperature maintained at 22.3 ± 1.0 °C and relative humidity at 57.6 ± 4.8% in both treatment rooms. Participants were randomly assigned to rooms to minimize environmental confounding.

The study evaluated the effects of both techniques using multiple criteria:

- Skin tension (firmness/stiffness) measured centrally on the thighs using EASYSTIFF® (standardized mapping of measurement sites),
- Cutaneous vascularization assessed by LAB-based colorimetric image analysis with standardized photographs before and after treatment,
- Patient-reported outcomes collected via a structured questionnaire.

All assessments were performed at baseline (before the first session) and after completion of the protocol, at identical anatomical sites on each thigh. Data collection and analysis were carried out by blinded operators.

This rigorous design allows for direct, intra-subject comparison of MMSA and traditional vacuum therapy effects, under strictly standardized environmental and procedural conditions.

### Statistical analysis

Statistical analysis was performed using R Studio. Data normality was assessed with the Shapiro– Wilk test. Homogeneity of variances was evaluated using Levene’s test. Depending on the distribution of the data, comparisons between treatment groups were made using either the Student’s t-test (for normally distributed data) or the Wilcoxon rank-sum test (for non-parametric data). Statistical significance was set at p < 0.05 (*), p < 0.005 (**), and p < 0.0005 (***).

## REFERENCES

Moortgat, P., Anthonissen, M., Meirte, J., Van Daele, U., & Maertens, K. (2016). The physical and physiological effects of vacuum massage on the different skin layers: a current status of the literature. Burns & Trauma, 4, 34. 10.1186/s41038-016-0053-9

Humbert, Ph., Fanian, F., Lihoreau, T., Jeudy, A., Elkhyat, A., Robin, S., et al. (2015). Mécano-Stimulation™ of the skin improves sagging score and induces beneficial functional modification of the fibroblasts: clinical, biological, and histological evaluations. Clinical Interventions in Aging, 10, 387–403. 10.2147/CIA.S69752

Palmieri, B., Aspiro, A., Sighinolfi, G., & Laurino, C. (2019). Intermittent microalveolar stromal stimulation therapy: lower legs functional and cosmetic reshaping by lympho-venous network activation. Minerva Chirurgica, 74 (Suppl. 1 N°3), 1–7.

Guo, Y., Song, Y., Xiong, S., Wang, T., Liu, W., Yu, Z., & Ma, X. (2022). Mechanical Stretch Induced Skin Regeneration: Molecular and Cellular Mechanism in Skin Soft Tissue Expansion. International Journal of Molecular Sciences, 23(17), 9622. 10.3390/ijms23179622

Li, Y., Li, L., Li, B., Liao, W., Yang, X., & Li, Y. (2023). Mechanical stretching induces fibroblast apoptosis through activating Piezo1 and then destroying actin cytoskeleton. International Journal of Medical Sciences, 20(6), 771–780. https://www.medsci.org/v20p0771.htm

Ogura, Y., Tanaka, Y., Hase, E., Yamashita, T., & Yasui, T. (2019). Texture analysis of second-harmonic-generation images for quantitative analysis of reticular dermal collagen fibre in vivo in human facial cheek skin. Experimental Dermatology, 28(8), 899–905. 10.1111/exd.13560

Dulińska-Molak, I., Pasikowska, M., Pogoda, K., Lewandowska, M., Eris, I., & Lekka, M. (2014). Age-related changes in the mechanical properties of human fibroblasts and its prospective reversal after anti-wrinkle tripeptide treatment. International Journal of Peptide Research and Therapeutics, 20(1), 77–85. 10.1007/s10989-013-9370-z

Timin, G. & Milinkovitch, M. C. (2023). High-resolution confocal and light-sheet imaging of collagen 3D network architecture in very large samples. iScience, 26(4), 106452. 10.1016/j.isci.2023.106452

Gonçalves, A. C., Guirro, R. R. J., Rossi, L. A., de Jesus Guirro, R., & de Oliveira, G. E. C. (2022). Effects of therapeutic ultrasound and paraffin with or without vacuum massage on biomechanical properties of grafted skin after burn: a randomized controlled trial. Revista da Associação Médica Brasileira, 68(12), 1628–1635. 10.1590/1806-9282.20220994

Fisher, G. J., Quan, T., Purohit, T., Shao, Y., Cho, M. K., He, T., et al. (2009). Collagen fragmentation promotes oxidative stress and elevates matrix metalloproteinase-1 in fibroblasts in aged human skin. The American Journal of Pathology, 174(1), 101–114. 10.2353/ajpath.2009.080599

Chlasta, J., et al. (2023). EASYSTIFF®, a portable and innovative device able to separately analyze each skin compartment for the evaluation of mechanical properties. Longdom, 11(3), 1–8. 10.1101/2023.07.13.548841

Qiao, N. et al. (2024). Contactless mechanical stimulation of the skin using shear waves. J Mech Behav Biomed Mater, 156, 106597. 10.1016/j.jmbbm.2024.106597

Wahlsten, A. et al. (2021). Mechanical stimulation induces rapid fibroblast proliferation and accelerates the early maturation of human skin substitutes. Biomaterials, 273, 120779. 10.1016/j.biomaterials.2021.120779

Mendes, N. et al. (2022). A narrative review of current striae treatments. Healthcare (Basel), 10(12), 2565. 10.3390/healthcare10122565

Papaccio, F. et al. (2022). Focus on the contribution of oxidative stress in skin aging. Antioxidants (Basel), 11(6), 1121. 10.3390/antiox11061121

Xia, C.Y. et al. (2014). Analysis of blood flow and local expression of angiogenesis-associated growth factors in infected wounds treated with negative pressure wound therapy. Mol Med Rep, 9(5), 1749–1754. 10.3892/mmr.2014.1997

Jaiswal, S. & Jawade, S. (2024). Microneedling in dermatology: A comprehensive review of applications, techniques, and outcomes. Cureus, 16(9), e70033. 10.7759/cureus.70033

Zerini, I. et al. (2015). Cellulite treatment: a comprehensive literature review. J Cosmet Dermatol, 14(3), 224–240. 10.1111/jocd.12154

Kołodziejczak, A., Adamiak, J., & Rotsztejn, H. (2025). Endermologie as a complementary therapy in medicine and surgery and an effective aesthetic procedure: A literature review. Applied Sciences, 15(8), 4313. 10.3390/app15084313

